# Differentiation-dependent telomeric long non-coding transcription in a model of skeletal myogenesis

**DOI:** 10.1101/000679

**Authors:** Scott W. Brouilette, Samir Ounzain, Vinit Sawhney, Kenta Yashiro, Yasunori Shintani, Kunihiko Takahashi, Steven R. Coppen, Takuya Narita, Kelli Torsney, Martin Carrier, Niall Campbell, Ken Suzuki

## Abstract

Telomeres comprise the distal ends of eukaryotic chromosomes, serve to maintain genomic integrity and are extended by the ribonucleoprotein telomerase. Recent evidence indicates that telomeres are transcribed to generate long non-coding RNAs (lncRNAs) and that these transcripts (TERRA) may inhibit telomerase activity. In this study we assessed telomerase activity and telomeric lncRNA expression in a mouse model of skeletal myogenesis. Using the C2C12 cell line we demonstrated decreased telomerase activity during differentiation into terminally-differentiated skeletal myotubes. Despite existing in a post-mitotic state, residual telomerase activity remained in C2C12 myotubes, indicating a role independent of telomere extension. Telomeric transcripts were detected in both myoblasts and myotubes, with reduced expression during differentiation correlating with reduced telomerase expression. Our data indicate that in a mouse model of skeletal myogenesis TERRA expression does not reduce telomerase activity, suggesting that their relationship is more complex than originally perceived; the role of telomeric derived lncRNAs in relation to telomerase activity may be cell-type specific. These findings raise the possibility for novel non-telomerase regulatory function for TERRA-lncRNAs during skeletal myogenesis.

## Introduction

Telomeres comprise the distal ends of eukaryotic chromosomes and consist of a highly conserved, repetitive sequence up to 150 kilobases (kb) in certain mouse strains (Kipling & Cooke 1990; Starling *et al*. 1990; Zijlmans *et al*. 1997). Although the full complement of telomeric functions remains to be determined, increasing evidence supports roles in maintenance of genomic integrity (Chan & Blackburn 2004). A region of telomeric DNA fails to be replicated during cell division, resulting in progressive telomere shortening. The ribonucleoprotein, telomerase, is capable of extending telomeres (Greider & Blackburn 1985), but while telomerase is generally active in germ and progenitor cells, its activity is repressed in human somatic and differentiated cells (Wright *et al*. 1996). In contrast, tissue specific expression has been reported in somatic/differentiated cells from mouse models (Prowse & Greider 1995). Recent reports indicate that rather than being inert structures, telomeric C-rich tracts are actively transcribed by RNA polymerase II (Azzalin *et al*. 2007), giving rise to telomere derived long non coding RNAs (lncRNAs) called TERRA. The phenomenon appears to be regulated by developmental status, telomere length, cellular stress and chromatin structure (Schoeftner & Blasco 2008). Furthermore, TERRA has been shown to block telomerase activity in both mouse embryonic stem cells, and the HeLa cell line (Schoeftner & Blasco 2008), raising the possibility that one function of TERRA is to provide a method of fine control of telomerase activity during growth and development. Additionally, TERRA may also act as a regulator of telomeric heterochromatin, analogous to other lncRNAs including HOTAIR and Xist (Luke & Lingner 2009).

Development of skeletal muscle is mediated by a coordinated series of events, beginning with commitment of mesodermal precursor cells to the skeletal muscle lineage followed by myoblast fusion and alterations in muscle-specific gene expression (Buckingham *et al*. 2003; Christ & Ordahl 1995; McKinsey *et al*. 2002), including MyoD for myoblast specification (Andres & Walsh 1996; Dedieu *et al*. 2002). Terminal differentiation is characterised by exit from the cell cycle, and telomerase down-regulation is considered to be an early, necessary step in the process. In light of this we set out to assess telomere, telomerase and TERRA lncRNA dynamics in a mouse model of skeletal myogenesis.

## Results

### Skeletal myogenesis

Reduction of serum concentration as cells approached confluence resulted in classic myoblast (MB) fusion followed by multi-nucleation by day 2 (2dMT), becoming more pronounced by day 4 (4dMT, Figure 1a). It has been reported that proliferating myoblasts exit the cell cycle when MyoD is at its highest level (Skapek *et al*. 1995) and consistent with this we observed the highest level of expression in proliferating MBs with a 48.5% reduction in expression at day 4 of differentiation (p<0.05, Figure 1b). During differentiation we also observed an 83% reduction (p<0.001) in the key cell cycle regulator, cyclin D1 at day 4 (Figure 1b). This down-regulation is critical for myogenesis as cyclin D1 is capable of inhibiting the ability of MyoD to transactivate muscle-specific genes (Skapek *et al*. 1995), including Mef2c (Dodou *et al*. 2003), transcription of which was consequently increased 148% (p<0.05) in 2dMT (Figure 1c). Myocyte stress 1 (MS1) is a recently identified downstream target of MyoD shown to play a key role in C2C12 differentiation (Ounzain *et al*. 2008). Consistent with this Ms1 transcription increased 129% in 2dMT (p<0.05) and 341% in 4dMT (p<0.05, Figure 1c). Finally, Creatine kinase M isozyme (CKM) transcript increased 123% in 4dMT (p<0.01), consistent with previous reports (Ito *et al*. 2005).

**Fig 1.**
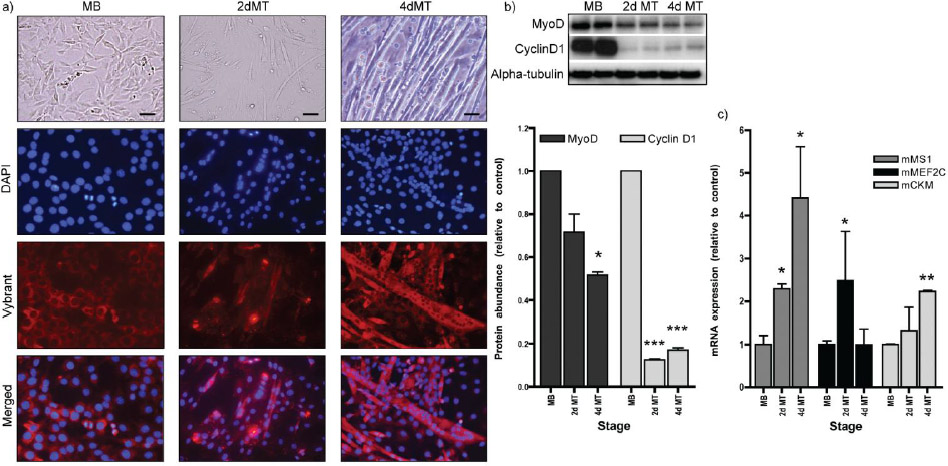
Skeletal myogenesis. **a)** C2C12 MBs showed classic morphology, beginning to form sporadic myotubes by 2dMT and becoming more pronounced by 4dMT. Vybrant staining of the cell membranes along with DAPI counter-staining of nuclei illustrated the transition from single nuclei myoblasts to multi-nuclear myotubes by day 4. Scale bar = 50μm. **b)** Western blotting demonstrated a progressive decrease in myoblast-specific MyoD expression during differentiation. The cell-cycle regulator Cyclin D1 was also significantly reduced as myoblasts differentiated into myotubes. **c)** RT-PCR analysis revealed increases in transcript levels of Ms1, Mef2c and Ckm, all with reported roles in skeletal myogenesis. * p<0.05, ** p<0.01, *** p<0.001 vs MB.

### Telomerase activity is decreased, but not absent, in C2C12 myotubes

Skeletal myogenesis is characterised by skeletal myoblast fusion to yield terminally-differentiated myotubes, while terminal differentiation itself is characterised by exit from the cell cycle. The ribo-nucleoprotein telomerase is capable of extending telomeres and expression is reported to occur during S-phase of the cell cycle (Zhao *et al*. 2008), thus activity is expected to diminish upon terminal differentiation. Figure 2a demonstrates telomerase activity in proliferating myotubes, with a 46% reduction in 2dMT and 59% in 4dMT (p<0.001 in both cases compared to MB). Thus, in contrast to previous reports (Bestilny *et al*. 1996; Holt *et al*. 1996) residual telomerase activity remained upon terminal differentiation.

**Fig 2.**
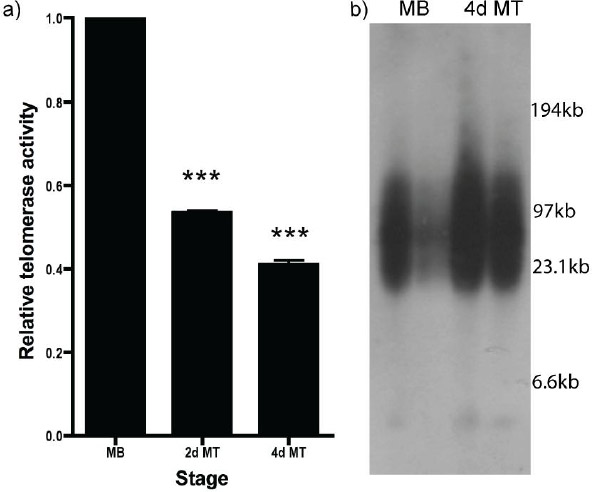
Telomerase and telomere dynamics during myogenesis. **a)** Telomerase PCR ELISA demonstrated that, relative to proliferating MBs, telomerase activity was significantly reduced, although not completely abrogated, in 2dMTs and 4dMTs. **b)**. Telomere length, measured by Pulse Field Gel Electrophoresis, showed no significant change during myogenesis. *** p<0.001 vs MB.

### Telomere length

Figure 2b demonstrates that telomere length, measured by pulse field gel electrophoresis, was similar between groups (25.4±0.17kb in MB and 26.92±0.58kb in 4dMT, p=0.1538).

### TERRA is expressed in both C2C12 myoblasts and myotubes

To assess proliferating and differentiated C2C12 cells for TERRA expression we used two independent methods. RNA-FISH revealed TERRA expression in 92.5±5.8% of proliferating MBs. This was significantly reduced to 40.7±13.2% in 2dMT and 35.8±13.5% in 4dMT (p<0.001 in both cases compared to MB, Figure 3a). Consistent with this, RNA dot-blotting indicated a 51% reduction in TERRA expression between proliferating MB and 4dMT (p=0.017, Figure 3b).

**Fig 3.**
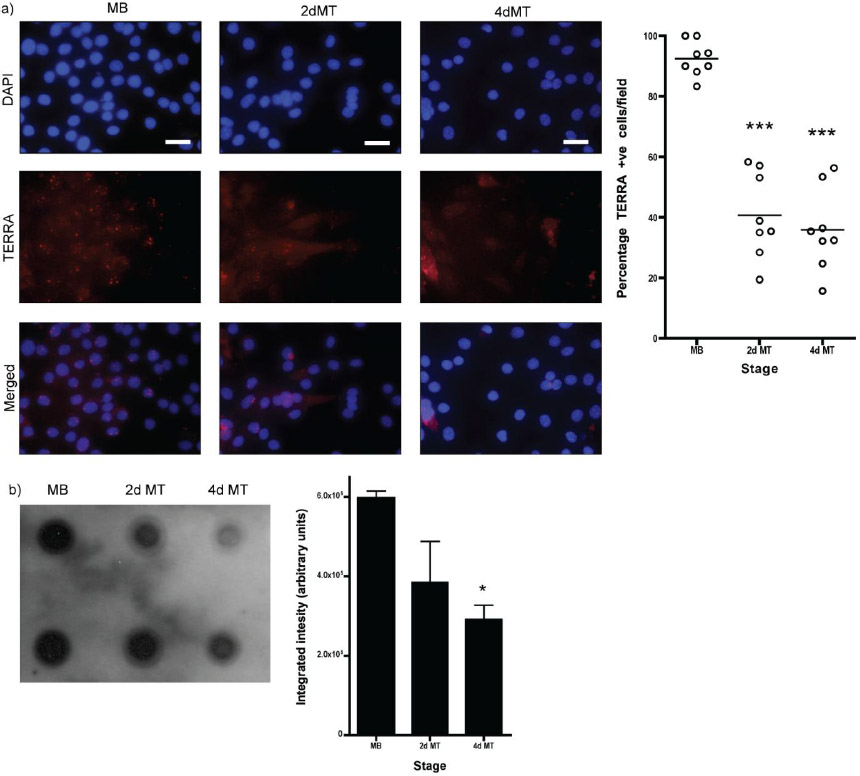
TERRA expression during myogenesis. **a)** RNA-FISH using a Cy-3 conjugated PNA probe and DAPI counter-stain revealed the presence of discrete TERRA foci in skeletal myoblasts, located to the nucleus. The frequency of cell nuclei exhibiting such foci was significantly less in 2dMTs and 4dMTs. **b)** RNA dot-blotting confirmed the decrease in TERRA during myogenesis. * p<0.05, *** p<0.001 vs myoblast. Scale bar = 50μm.

### Abrogation of telomerase activity by synthetic TERRA

TERRA possesses complimentarity to the RNA template of the telomerase complex, and has been shown to inhibit telomerase activity in the TRAP assay (Schoeftner & Blasco 2008). To assess the impact of TERRA on telomerase activity in our model protein extracts were incubated with increasing amounts of the synthetic TERRA oligonucleotide, T1 ([UUAGGG]_3_), prior to the assay. Addition of T1 resulted in a significant reduction in telomerase activity at all concentrations tested (over 90%, p<0.01 in all cases, Figure 4).

**Fig 4.**
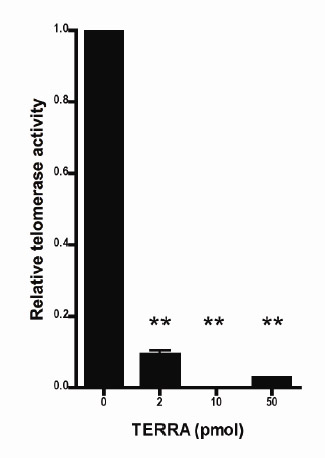
Telomerase inhibition by TERRA. Addition of synthetic TERRA inhibited telomerase activity, assessed by TRAP PCR-ELISA, at all concentrations tested. Activity was reduced in excess of 90%. ** p<0.01 vs control (no TERRA).

## Discussion

In the present study we have characterised the changes in telomerase activity and TERRA lncRNA expression that occur during the differentiation of the mouse C2C12 cell line from proliferating myoblasts to terminally-differentiated myotubes. In contrast to reports indicating almost complete abrogation of telomerase activity during myogenesis we report only a 50% reduction, indicating a role independent of telomere length maintenance in post-mitotic cells. Furthermore, telomeric transcripts (TERRA) were observed in both C2C12 myoblasts and myotubes, with reduced expression during myogensis correlating with reduced telomerase activity. Given recent reports indicating that increased TERRA expression down-regulates telomerase activity (Ng *et al*. 2009; Schoeftner & Blasco 2008), our novel observation implies a complex relationship between TERRA and telomerase activity during skeletal myogenesis.

Our findings expand on the limited data relating to telomerase down-regulation in terminal differentiation during skeletal myogenesis. Unlike previous reports that indicate a greater than 90% down-regulation as the cells exit the cell cycle and differentiate into myotubes (Bestilny *et al*. 1996; Holt *et al*. 1996), we observed residual telomerase activity at relatively high levels (50% of the maximum in highly proliferative myoblasts). This may reflect technical differences between the methodologies, with the TRAP-ELISA reporting levels of activity that were previously below the threshold of detection using the original assay. While telomerase was originally thought to be superfluous in cells that had exited the cell cycle, there is recent evidence suggesting that telomerase may have functions that are independent of telomere length maintenance. During oxidative stress telomerase has been shown to be actively exported from the nucleus to the cytoplasm where it co-localises with mitochondria (Ahmed *et al*. 2008), preserving function by binding to mitochondrial DNA and protecting against oxidative stress-induced damage (Haendeler *et al*. 2009). Proteomic analysis of C2C12 cells indicates that respiratory chain proteins are up-regulated by 10 days post-differentiation (Kislinger *et al*. 2005), resulting in increased potential for generation of oxidative stress. This is countered, in part, by up-regulation of uncoupling protein-3 (Flandin *et al*. 2005). Given that these data illustrate the importance of managing the increased oxidative stress that can be generated by differentiated skeletal muscle, our data indicating residual telomerase activity in post-mitotic cells suggests a potential role for telomerase, independent of its role in maintaining telomeric structure. This may represent a pre-emptive, protective mechanism, as the cell prepares itself to cope with the increased oxidative stress that may ensue as myotube formation is completed and muscle contraction begins.

The current model for TERRA regulation of telomerase activity proposes that TERRA anneals to the RNA sub-unit of telomerase, abrogating its use as a template for telomere extension (Schoeftner & Blasco 2008), however our data demonstrating a significant reduction in TERRA expression during differentiation, concurrent with reduced telomerase activity, suggest an alternative relationship during myogenesis. While we report a positive correlation between TERRA and telomerase activity in the cell model we did observe a negative correlation in an *in vitro* assay of telomerase activity, indicating that TERRA does indeed inhibit telomerase activity when simply incubated with protein extracts from the C2C12 cell line. One potential explanation for our observed changes in TERRA during C2C12 differentiation relates to SMG proteins, several of which are involved in telomere maintenance. Upf1 in particular is up-regulated over 5-fold during myogeneis in C2C12 cells (Kim *et al*. 2007), and Azzalin *et al* (Azzalin *et al*. 2007) have shown that Upf1 promotes the displacement of TERRA from telomeric chromatin. TERRA appears to be under tight transcriptional control (Schoeftner & Blasco 2008); while telomere-associated TERRA may promote genomic stability (Horard & Gilson 2008), detachment likely leads to degradation. This model would be consistent with our data: telomeric transcription may remain high in myotubes (or may actually increase in an attempt to fine-tune down-regulation of telomerase activity), but the increased Upf1 reported in C2C12 myotubes may displace TERRA resulting in rapid degradation, thus we observed a reduced signal.

A second mechanism involving methylation may also play a role. Methylation is an epigenetic mechanism that generally results in silencing of the associated sequence. In humans sub-telomeric DNA is hypo-methylated in sperm and ova, and is subjected to heavy *de novo* methylation during development (Brock *et al*. 1999; de Lange *et al*. 1990). Recently it has been demonstrated that disease-associated hypo-methylation of the sub-telomere in humans is accompanied by increased TERRA (Yehezkel *et al*. 2008); rather than a primary role in telomerase regulation, TERRA formation may reflect altered epigenetic status at the sub-telomeric region. Importantly, the two mechanisms outlined are not mutually exclusive. Finally, it is also possible that during myogenic differentiation TERRA mediates an exclusively non-telomerase/telomeric functional role. Considering the increasingly vast repertoire of regulatory functions encoded by long non coding RNAs we suspect TERRA function could be diverse and cell-type/context specific (Wang & Chang 2011; Wapinski & Chang 2011). To conclude, we provide the first report of telomeric lncRNA expression in a mouse model of skeletal myogenesis, and demonstrate a positive association with telomerase activity. These data are in contrast to previous reports, albeit using different cell types, suggesting that TERRA inhibits telomerase activity, and indicate that the relationship may be cell-type specific and, thus, more complex than originally believed. Importantly, this study raises several questions regarding the relationship between TERRA expression and telomerase activity during skeletal myogenesis.

## Experimental Procedures

### Cell culture

C2C12 mouse skeletal myoblasts were cultured in DMEM (Invitrogen) supplemented with 20% fetal calf serum, 2mM glutamine, penicillin (100 units/ml, Sigma) and streptomycin (100μg/ml, Sigma) and maintained at 37°C in 5% CO_2_. Once cells exceeded 90% confluence media was replaced with DMEM supplemented with 2% horse serum to induce differentiation. Differentiation media was changed every 2 days.

### Morphology and myotube formation

For assessment of myogenic differentiation, C2C12 cells were cultured in chamber slides and examined as proliferating myoblasts (MB), 2 day (2dMT) and 4 day myotubes (4dMT). For assessment of cellular fusion media was aspirated and replaced with fresh media containing Vybrant Cell Labeling Solution (Invitrogen) at 1:200 dilution and incubated at 37°C for 20 minutes, followed by three 10 minute washes in fresh, unlabeled media. After washing in phosphate-buffered saline (PBS), cells were fixed in 4% paraformaldehyde for 10 minutes, washed in 70% ethanol and mounted (Dako Fluorescent Mount containing 200ng/ml 4’,6-diamidino-2-phenylindole (DAPI) counter-stain). Images were captured on a Keyance All-In-One Fluorescent microscope (Keyance, Japan).

### Western blotting

Total protein was isolated using SB_20_ buffer (50mM Tris pH 7.4, 5mM EDTA, 20% SDS), quantitated using BioRad D_C_ Protein Assay (BioRad), electrophoresed in SDS-polyacrylamide gels (Invitrogen) and transferred to Hybond ECL (GE Healthcare).

Membranes were probed with antibodies against MyoD (Santa Cruz) and Cyclin D1 (Millipore), normalized to α-tubulin (Sigma). Autoradiographs were scanned and image analysis performed using ImageJ (ImageJ: http://rsb.info.nih.gov/ij).

### RNA isolation

Cells were trypsinized and total RNA was isolated using the RNeasy Mini Kit (QIAgen) according to manufacturer’s guidelines. On-column DNase I digestion (RNase-Free DNase Set, QIAgen) was performed to eliminate genomic DNA contamination.

### RT-PCR

Total RNA (2μg) was reverse transcribed to cDNA using oligo (dT) and Superscript II Reverse Transcriptase (Invitrogen) according to the manufacturers instructions. Quantitative PCR (QPCR) was executed using a 7500 Real-Time PCR System (Applied Biosystems).

Briefly, a master-mix containing (per reaction) 12.5μl SYBR-Green Jump Start Taq Readymix (Sigma-Aldrich) and 5μl oligonucleotides (5pmol of forward and reverse primers) were aliquoted into Optical 96-well reaction plates (Applied Biosystems), and the cDNA template added (7.5μl, 1/15 dilution in ddH2O). PCR cycling parameters for all primer pairs were 50°C for 2 minutes, 95°C for 10 minutes, followed by 40 cycles of 95°C for 15 seconds and 59°C for 60 seconds. Subsequent to QPCR, a dissociation curve analysis was performed to check for aberrant amplification products. QPCR analysis of Gapdh was performed in each 96-well plate as an endogenous control and the relative quantitation protocol was used.

Primers for QPCR were as follows: Gapdh: forward 5’-ACC ACA GTC CAT GCC ATC AC-3’, reverse 5’-TCC ACC ACC CTG TTG CTG TA-3’; Ckm: forward 5’-TCA AGG GTT ACA CTC TGC CT-3’, reverse 5’-TTC ACC CAC ACA AGG AAG C-3’; Mef2c: forward 5’-ATT CCT GCT GTT CCA CCT CC-3’, reverse 5’-AAC GCG GAG ATC TGG CTT AC-3’; Ms1: forward 5’-GTG ACA GCA TAG ACA CAG GAG GAC-3’. reverse 5’-CAC TGC TGC CCA CCT GCC TT-3’. All samples run in duplicate.

### Telomerase Activity

Telomerase activity was assessed using TeloTAGGG Telomerase PCR ELISA (Roche) according to manufacturer’s instructions. Briefly, at each time-point 2×10^5^ cells were washed in PBS, pelleted and lysed to release protein. Samples were assayed in duplicate, using 5000 cell equivalents per PCR reaction, and 5μl of PCR product per ELISA. HeLa cells were used as a positive control; heat inactivated samples (10 minutes at 85°C prior to PCR) served as negative controls.

### Telomere Length

Telomere length was assessed using pulse-field gel electrophoresis (PFGE). Briefly, at each time-point cells were harvested and 10^7^ cells embedded in agarose gel plugs (CHEF Genomic DNA Plug Kit, BioRad) and digested with Rsa1/Hinf1 (Invitrogen). Gel plug pieces containing approximately 2×10^6^ cells were subjected to PFGE, transferred to Hybond N^+^ (GE Healthcare) and telomeric signals visualized using TeloTAGGG Telomere Length Assay (Roche). Autoradiographs were scanned and image analysis performed using ImageQuant (Molecular Dynamics), as previously described (Brouilette *et al*. 2003; Vasa-Nicotera *et al*.2005).

### RNA-Fluorescence in situ hybridization (FISH)

RNA-FISH was performed as previously described (Azzalin *et al*. 2007). Briefly, cells were cultured in chamber-slides (Nunc) until reaching required degree of confluency/differentiation. Slides were washed in PBS and incubated for 7 minutes in ice-cold cyto-skeletal (CSK) Buffer (100mM NaCl, 300mM sucrose, 3mM MgCl2, 10mM PIPES pH 7.0, 0.5% Triton-X) containing either 10mM Vanadyl Ribonucleoside Complex (New England Biolabs) or 1mg/ml Rnase A (Invitrogen). Slides were washed again in PBS, fixed in 4% paraformaldehyde for 10 minutes at room temperature, and dehydrated in an ethanol series (70%, 90% and 100%, 5 minutes each). After air-drying slides were hybridised (2x SSC, 2mg/ml BSA, 10% dextran sulphate, 20% formamide, 2.0ug/ml PNA-telomere probe, 10mM Vanadyl Ribonucleoside Complex) overnight at 37°C. Slides were then washed twice with 2X SSC/50% formamide at 37°C, twice with 2X SSC at 37°C, and once with 2X SSC at room temperature. Slides were mounted with fluorescent mounting media (Dako) and DAPI counterstain (200ng/ml). Images were captured using a Keyance All-In-One Fluorescent microscope (Keyance, Japan) using 40x magnification and identical settings for both myoblast and myotube slides (9 second exposure time in Cy-5 channel for TERRA). TERRA formation was quantitated as percentage of TERRA positive DAPI-stained nuclei per field (7–10 random fields per time-point).

### RNA dot-blot

Total RNA (5μg) was spotted in duplicate onto Hybond N^+^ (GE Healthcare) with RNase A treated samples included as controls. Blots were air-dried, UV cross-linked using the Autocrosslink Mode (Stratagene) and pre-hybridised for 1 hour at 55°C in Easy Hyb solution (Roche). 100pmol telomere probe (CCCTAA)_3_ was labeled with DIG using DIG Oligonucleotide 3’-End-Labeling Kit (Roche) and hybridised overnight at 55°C. Following low stringency (2xSSC/0.1% SDS at room temperature) and high stringency (0.2xSSC/0.1% SDS at 55°C) washes, blots were visualised using the DIG Northern Starter Kit (Roche).

Blots were stripped and re-probed with Gapdh as control. Autoradiographs were scanned and image analysis performed using ImageJ (ImageJ: http://rsb.info.nih.gov/ij).

### Synthetic TERRA

Protein lysates (5000 cell equivalents) from C2C12 myoblasts were incubated with increasing amounts (2, 10 and 50pmoles) of T1 ([UUAGGG]_3_) oligonucleotide (50μl total reaction volume) prior to the telomerase extension reaction in Telomerase PCR ELISA.

### Statistical Analysis

Differences in percentage of TERRA foci per field (FISH), TERRA expression (RNA dot-blot) and telomerase activity (TRAP-ELISA) at each time point were compared by one-way ANOVA and Bonferroni’s Multiple Comparison Test. A p-value <0.05 was considered significant.

All analyses were performed using GraphPad Prism software (Version 4, 2005).

## Acknowledgements

This work was supported by the Barts and The London Charity.

## References

Ahmed, S., Passos, J.F., Birket, M.J., et al. (2008) Telomerase does not counteract telomere shortening but protects mitochondrial function under oxidative stress. Journal of cell science 121, 1046–1053.

Andres, V. & Walsh, K. (1996) Myogenin expression, cell cycle withdrawal, and phenotypic differentiation are temporally separable events that precede cell fusion upon myogenesis. The Journal of cell biology 132, 657–666.

Azzalin, C.M., Reichenbach, P., Khoriauli, L., Giulotto, E. & Lingner, J. (2007) Telomeric repeat containing RNA and RNA surveillance factors at mammalian chromosome ends. Science 318, 798–801.

Bestilny, L.J., Brown, C.B., Miura, Y., Robertson, L.D. & Riabowol, K.T. (1996) Selective inhibition of telomerase activity during terminal differentiation of immortal cell lines. Cancer research 56, 3796–3802.

Brock, G.J., Charlton, J. & Bird, A. (1999) Densely methylated sequences that are preferentially localized at telomere-proximal regions of human chromosomes. Gene 240, 269–277.

Brouilette, S., Singh, R.K., Thompson, J.R., Goodall, A.H. & Samani, N.J. (2003) White cell telomere length and risk of premature myocardial infarction. Arteriosclerosis, thrombosis, and vascular biology 23, 842–846.

Buckingham, M., Bajard, L., Chang, T., et al. (2003) The formation of skeletal muscle: from somite to limb. Journal of anatomy 202, 59–68.

Chan, S.R. & Blackburn, E.H. (2004) Telomeres and telomerase. Philosophical transactions of the Royal Society of London. Series B, Biological sciences 359, 109–121.

Christ, B. & Ordahl, C.P. (1995) Early stages of chick somite development. Anatomy and embryology 191, 381–396.

de Lange, T., Shiue, L., Myers, R.M., et al. (1990) Structure and variability of human chromosome ends. Molecular and cellular biology 10, 518–527.

Dedieu, S., Mazeres, G., Cottin, P. & Brustis, J.J. (2002) Involvement of myogenic regulator factors during fusion in the cell line C2C12. The International journal of developmental biology 46, 235–241.

Dodou, E., Xu, S.M. & Black, B.L. (2003) mef2c is activated directly by myogenic basic helix-loop-helix proteins during skeletal muscle development in vivo. Mechanisms of development 120, 1021–1032.

Flandin, P., Donati, Y., Barazzone-Argiroffo, C. & Muzzin, P. (2005) Hyperoxia-mediated oxidative stress increases expression of UCP3 mRNA and protein in skeletal muscle. FEBS letters 579, 3411–3415.

Greider, C.W. & Blackburn, E.H. (1985) Identification of a specific telomere terminal transferase activity in Tetrahymena extracts. Cell 43, 405–413.

Haendeler, J., Drose, S., Buchner, N., et al. (2009) Mitochondrial telomerase reverse transcriptase binds to and protects mitochondrial DNA and function from damage. Arteriosclerosis, thrombosis, and vascular biology 29, 929–935.

Holt, S.E., Wright, W.E. & Shay, J.W. (1996) Regulation of telomerase activity in immortal cell lines. Molecular and cellular biology 16, 2932–2939.

Horard, B. & Gilson, E. (2008) Telomeric RNA enters the game. Nature cell biology 10, 113– 115.

Ito, H., Ueda, H., Iwamoto, I., et al. (2005) Nordihydroguaiaretic acid (NDGA) blocks the differentiation of C2C12 myoblast cells. Journal of cellular physiology 202, 874–879.

Kim, Y.K., Furic, L., Parisien, M., Major, F., DesGroseillers, L. & Maquat, L.E. (2007) Staufen1 regulates diverse classes of mammalian transcripts. The EMBO journal 26, 2670– 2681.

Kipling, D. & Cooke, H.J. (1990) Hypervariable ultra-long telomeres in mice. Nature 347, 400–402.

Kislinger, T., Gramolini, A.O., Pan, Y., Rahman, K., MacLennan, D.H. & Emili, A. (2005) Proteome dynamics during C2C12 myoblast differentiation. Molecular & cellular proteomics: MCP 4, 887–901.

Luke, B. & Lingner, J. (2009) TERRA: telomeric repeat-containing RNA. The EMBO journal 28, 2503–2510.

McKinsey, T.A., Zhang, C.L. & Olson, E.N. (2002) Signaling chromatin to make muscle. Current opinion in cell biology 14, 763–772.

Ng, L.J., Cropley, J.E., Pickett, H.A., Reddel, R.R. & Suter, C.M. (2009) Telomerase activity is associated with an increase in DNA methylation at the proximal subtelomere and a reduction in telomeric transcription. Nucleic acids research 37, 1152–1159.

Ounzain, S., Dacwag, C.S., Samani, N.J., Imbalzano, A.N. & Chong, N.W. (2008) Comparative in silico analysis identifies bona fide MyoD binding sites within the Myocyte stress 1 gene promoter. BMC molecular biology 9, 50.

Prowse, K.R. & Greider, C.W. (1995) Developmental and tissue-specific regulation of mouse telomerase and telomere length. Proceedings of the National Academy of Sciences of the United States of America 92, 4818–4822.

Schoeftner, S. & Blasco, M.A. (2008) Developmentally regulated transcription of mammalian telomeres by DNA-dependent RNA polymerase II. Nature cell biology 10, 228– 236.

Skapek, S.X., Rhee, J., Spicer, D.B. & Lassar, A.B. (1995) Inhibition of myogenic differentiation in proliferating myoblasts by cyclin D1-dependent kinase. Science 267, 1022– 1024.

Starling, J.A., Maule, J., Hastie, N.D. & Allshire, R.C. (1990) Extensive telomere repeat arrays in mouse are hypervariable. Nucleic acids research 18, 6881–6888.

Vasa-Nicotera, M., Brouilette, S., Mangino, M., et al. (2005) Mapping of a major locus that determines telomere length in humans. American journal of human genetics 76, 147–151.

Wang, K.C. & Chang, H.Y. (2011) Molecular mechanisms of long noncoding RNAs. Molecular cell 43, 904–914.

Wapinski, O. & Chang, H.Y. (2011) Long noncoding RNAs and human disease. Trends in cell biology 21, 354–361.

Wright, W.E., Piatyszek, M.A., Rainey, W.E., Byrd, W. & Shay, J.W. (1996) Telomerase activity in human germline and embryonic tissues and cells. Developmental genetics 18, 173– 179.

Yehezkel, S., Segev, Y., Viegas-Pequignot, E., Skorecki, K. & Selig, S. (2008) Hypomethylation of subtelomeric regions in ICF syndrome is associated with abnormally short telomeres and enhanced transcription from telomeric regions. Human molecular genetics 17, 2776–2789.

Zhao, Y.M., Li, J.Y., Lan, J.P., et al. (2008) Cell cycle dependent telomere regulation by telomerase in human bone marrow mesenchymal stem cells. Biochemical and biophysical research communications 369, 1114–1119.

Zijlmans, J.M., Martens, U.M., Poon, S.S., et al. (1997) Telomeres in the mouse have large inter-chromosomal variations in the number of T2AG3 repeats. Proceedings of the National Academy of Sciences of the United States of America 94, 7423–7428.

